# The Homo-Edit Distance Problem

**DOI:** 10.1101/2020.05.27.118273

**Authors:** Maren Brand, Nguyen Khoa Tran, Philipp Spohr, Sven Schrinner, Gunnar W. Klau

**Author notes:** Shared first authors (Maren Brand, Nguyen Khoa Tran).

## Abstract

We consider the homo-edit distance problem, which is the minimum number of homo-deletions or homo-insertions to convert one string into another. A homo-insertion is the insertion of a string of equal characters into another string, while a homo-deletion is the inverse operation. We show how to compute the homo-edit distance of two strings in polynomial time: We first demonstrate that the problem is equivalent to computing a common subsequence of the two input strings with a minimum number of homo-deletions and then present a dynamic programming solution for the reformulated problem.

**2012 ACM Subject Classification:** Applied computing → Bioinformatics; Applied computing → Molecular sequence analysis; Theory of computation → Dynamic programming

## 1 Introduction

A homo-insertion is an insertion of a string of equal characters, which we also call a *block*, into another string. A homo-deletion is the inverse operation, that is, the deletion of such a block. We consider the following problem: Given two strings, what is the minimum number of homo-insertions or homo-deletions needed to convert one into the other? We refer to this number as the homo-edit distance. This distance is a generalization of the edit distance between two strings, where only insertions and deletions are possible, which is also known as the longest common subsequence distance [4, 1]. Unlike in the classic special case, where blocks consist only of single characters, two blocks may merge to one after a homo-deletion. For example, the homo-edit distance of `ATA` and the empty string is 2 and is achieved by first deleting `T` and then the block `AA`. This property makes the homo-edit distance more difficult to compute than the longest common subsequence distance.

We became aware of this problem as an exercise (6.40) in the classic textbook on bioinformatics algorithms by Jones and Pevzner [3]. As this exercise caused our students a lot of trouble we decided to look at it more closely within a thesis project [2]. We show how to compute the homo-edit distance of two strings in polynomial time: We first demonstrate that the problem is equivalent to computing a common subsequence of the two input strings with a minimum number of homo-deletions and then present a dynamic programming solution for the reformulated problem.

## 2 Problem Formulation

Let Σ be a finite alphabet. A string of length *n* ∈ ℕ_0_ is defined as *s* = *s*_1_*s*_2_ … *s*_*n*_ ∈ Σ^*n*^. The empty string is denoted as *ε*. We also write the length of a string *s* as |*s*|. A *block* is a string consisting of identical characters, and we write *a*^*k*^ for a block of length *k* and some *a* ∈ Σ. We refer to substrings of *s* by

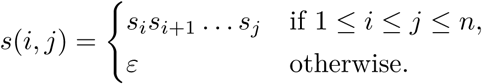

Subsequences 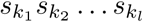 of *s* are characterized by their indices *k*_1_, *k*_2_, *…, k*_*l*_, where 1 ≤ *k*_1_ *< k*_2_ *< … < k*_*l*_ ≤ *n*.

We define two string operations which we subsume as *homo-operations*: The first operation inserts a block of length *k* into a string at a certain position. Let *a* ∈ Σ and let *u* = *a*^*k*^ be the block that is to be inserted into string *s* at position *i*, where 1 ≤ *i* ≤ *n* + 1. We define this *homo-insertion* as the string

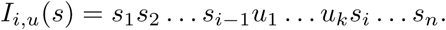

The second operation deletes a block *s*(*i, j*) = *a … a* with *a* ∈ Σ and 1≤ *i* ≤ *j* ≤ *n*. We define this *homo-deletion* as the string

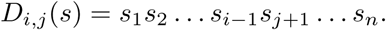

Note that both operations are reversible, that is, for each homo-insertion there is a homo-deletion that can be applied to obtain the original string, and vice versa. For an operation 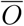 we denote the corresponding reverse operation by *O*. Reversibility also holds for chains of operations as the following lemma shows.

### Lemma 1.

*Consider a series of homo-operations O*_1_, *O*_2_, *…, O*_*k*_ *to convert a string s into another string t, that is, t* = (*O*_*k*_ ○ *O*_*k*−1_ ○ *…* ○ *O*_1_)(*s*). *Then, also* 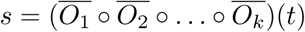 *holds.*

**Proof by induction.**

▪ Base case: *k* = 1 Case 1: *t* = *O*_1_(*s*) = *I*_*i,u*_(*s*) is obtained by a homo-insertion of a string *u* = *a*^*j*−*i*+1^ into *s* at position *i*, where *a* ∈ Σ and *j* ≥ *i*. We reverse this homo-insertion by using a homo-deletion of substring *s*(*i, j*) = *u* from *t*, i.e., 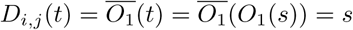. Case 2: *t* = *O*_1_(*s*) = *D*_*i,j*_(*s*) is obtained by a homo-deletion of a substring *u* = *s*(*i, j*) from *s*. We reverse this homo-deletion by using a homo-insertion of *u* into *t* at position *i*, i.e., 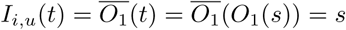.
▪ Induction step: *k* → *k* + 1 Let *s*′ = (*O*_*k*_ ○ *O*_*k*−1_ ○ *…* ○ *O*_1_)(*s*) such that *O*_*k*+1_(*s*′) = *t*. Case 1: *t* = *O*_*k*+1_(*s*′) = *I*_*i,u*_(*s*′) is obtained by a homo-insertion of a string *u* = *a*^*j*−*i*^ into *s*′at position *i*, where *a* ∈ Σ and *j* ≥ *i*. We reverse this homo-insertion by using a homo-deletion of substring *s*′(*i, j*) = *u* from *t*, i.e., 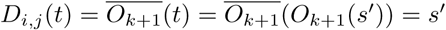. Case 2: *t* = *O*_*k*+1_(*s*) = *D*_*i,j*_(*s*′) is obtained by a homo-deletion of a substring *u* = *s*′ (*i, j*) from *s*′. We reverse this homo-deletion by using a homo-insertion of *u* into *t* at position *i*, i.e., 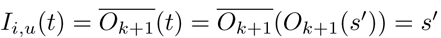. ◂

We define the *homo-edit distance H*(*s, t*) between two strings *s* and *t* as the minimum number of homo-operations to convert *s* into *t*. From Lemma 1 it follows that the homo-edit distance is symmetric, that is, *H*(*s, t*) = *H*(*t, s*). We can now define the homo-edit distance problem formally as follows:

### Problem 1

(Homo-Edit Distance Problem). Given two strings *s* and *t*, compute their homo-edit distance *H*(*s, t*).

## 3 Problem Reformulation

In this section we point out that the homo-edit distance between two strings *s* and *t* can be computed by considering homo-deletions only. For this we show that there exists a common subsequence *v* of both strings such that converting both *s* and *t* into *v* needs a total of *H*(*s, t*) homo-deletions.

### Lemma 2.

*Let s and t be two strings and let H*(*s, t*) = *k. Then there exists an optimal series of homo-operations O*_1_, *O*_2_, *…, O*_*k*_ *to convert s into t, such that the first part of the series contains only homo-deletions and the second part only homo-insertions.*

**Proof.**

Let 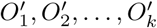 be any optimal series of homo-operations without the property stated in Lemma 2, let 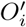 be a homo-insertion followed by a homo-deletion 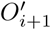, where 1 ≤ *i* ≤ *k* − 1, and let 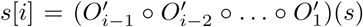. If there exists a homo-deletion 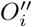 followed by a homo-insertion 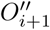 such that 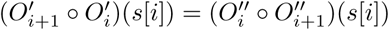, then the series *O*_1_, *O*_2_, *…, O*_*k*_ must exist as well, because we can repeatedly replace each homo-insertion followed by a homo-deletion with a homo-deletion followed by a homo-insertion, resulting in the same string.

Let *u* be the string that we want to insert by applying *O*_*i*_ and let *w* be the string that we want to delete by applying 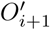. We consider two cases:

▪ Case 1: *w* consists of a substring of *s*[*i*] only. Let *p*_1_ be the position in *s*[*i*] where we want to insert *u*, and let *p*_2_ be the position of *w*_1_ in *s*[*i*]. We can safely delete *w* first and then insert *u* either at position *p*_1_, if *p*_1_ ≤ *p*_2_, or at position *p*_1_ − |*w*|, if *p*_1_ *> p*_2_.
▪ Case 2: *w* consists of a substring of *u* as well as a substring of *s*[*i*]. Let *a* ∈ Σ, let 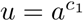, and let 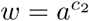, such that after applying both homo-operations, we either inserted or deleted 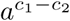, depending on whether *c*_1_ *> c*_2_ or *c*_1_≤ *c*_2_. This means we could use one instead of two homo-operations for inserting or deleting 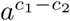, or even zero if *c*_1_ = *c*_2_. Thus, the series 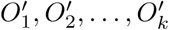 would not be optimal, which is a contradiction. ◂

### Lemma 3.

*Let s and t be two strings. Then H*(*s, t*) = *k if and only if there is a common subsequence v of s and t, such that it takes a total of k homo-deletions to convert s and t into v.*

**Proof.**

From Lemma 2 we know that for converting *s* into *t*, there exists a series of homo-operations *O*_1_, *O*_2_, … *O*_*k*_ where the first *i* homo-operations of this series include homo-deletions only and where the last *k* − *i* homo-operations include homo-insertions only, where 0 ≤ *i* ≤ *k*. Let *v* be the string that we obtain by performing these homo-deletions on *s*, that is, *v* = (*O*_*i*_ ○ *O*_*i*−1_ ○ *…* ○ *O*_1_)(*s*). Then *v* is a subsequence of *s* by definition. From Lemma 1 we know that we can reverse the homo-insertions of the series *t* = (*O*_*k*_ ○ *O*_*k*−1_ ○ *…* ○ *O*_*i*+1_)(*v*) such that 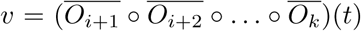. Thus, *v* is also a subsequence of *t*, and we can obtain *v* by a total of *k* homo-deletions. ◂

Lemma 3 implies that we can safely disregard homo-insertions for computing homo-edit distances. In the next section we present an algorithm that computes the homo-edit distance of two strings by finding the minimum number of homo-deletions to convert both into a common subsequence.

## 4 Dynamic Programming Algorithm

This section contains our algorithmic contributions to the problem, their correctness proofs, a note on backtracking and a running time analysis.

### 4.1 Algorithms

We compute the homo-edit distance between two strings *s* and *t* with a two-part dynamic programming (DP) algorithm: The first part is a precomputation step that computes and stores the homo-edit distance between every substring of both *s* and *t* and the empty string *ε*. The second part is the main algorithm that, similar to classic textbook approaches for sequence alignment, computes a DP matrix containing the homo-edit distances between all prefixes of *s* and *t*. For better understanding we explain the main algorithm first.

Given two strings *s* and *t*, let *v* be an optimal common subsequence, that is, *v* satisfies the conditions of Lemma 3. Let *m* = |*s*| and *n* = |*t*|. We compute an (*m* + 1) × (*n* + 1) matrix *d*, where each entry *d*_*i,j*_ corresponds to the homo-edit distance between the prefixes *s*(1, *i*) and *t*(1, *j*), with the following recurrence:

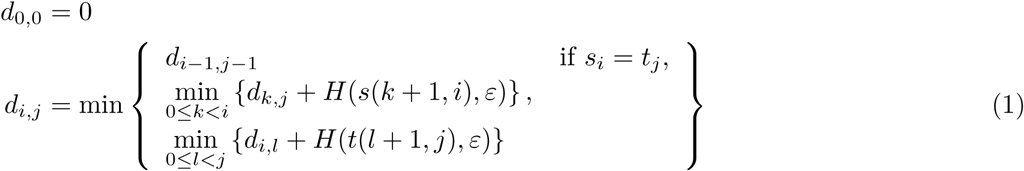

We start by initializing *d*_0,0_ with 0. For all other entries we proceed, e.g., from top to bottom (*i* = 0, 1, *…, m*) and from left to right (*j* = 0, 1, *…, n*), and consider three cases for the homo-edit distance between *s*(1, *i*) and *t*(1, *j*), among which we pick the minimum:

1. The first case is given if we have a match, i.e., *s*_*i*_ = *t*_*j*_. In this case, the common character could be part of an optimal common subsequence *v*. As we would neither delete *s*_*i*_ nor *t*_*j*_ by a homo-deletion, we have *d*_*i,j*_ = *d*_*i*−1,*j*−1_.
2. The next case comprises all possibilities that involve deleting *s*_*i*_ from *s*, meaning that this character would not be part of an optimal subsequence *v*. More precisely, for *d*_*i,j*_ we consider each entry *d*_*k,j*_ of the same column *j* in a row *k* from above plus the cost of deleting *s*(*k* + 1, *i*). We will show how to compute the homo-edit distances between all substrings of a string and the empty string *ε* later.
3. The last case consists of all possibilities where we delete *t*_*j*_ from *t*. More precisely, for *d*_*i,j*_ we consider each entry *d*_*i,l*_ of the same row *i* in a column *l* from left plus the cost of deleting *t*(*l* + 1, *j*).

Eventually, *d*_*m,n*_ contains the homo-edit distance between *s* and *t*. We can obtain an optimal subsequence *v* and thereby an optimal series of operations to obtain *s* from *t* or vice versa by backtracking the cases from *d*_*m,n*_ to *d*_0,0_. Note that there can be multiple possibilities for *v*. See Algorithm 1 and the paragraph about backtracking below for more details.

#### Algorithm 1 Main dynamic programming algorithm to compute the homo-edit distance between two strings *s* and *t*.

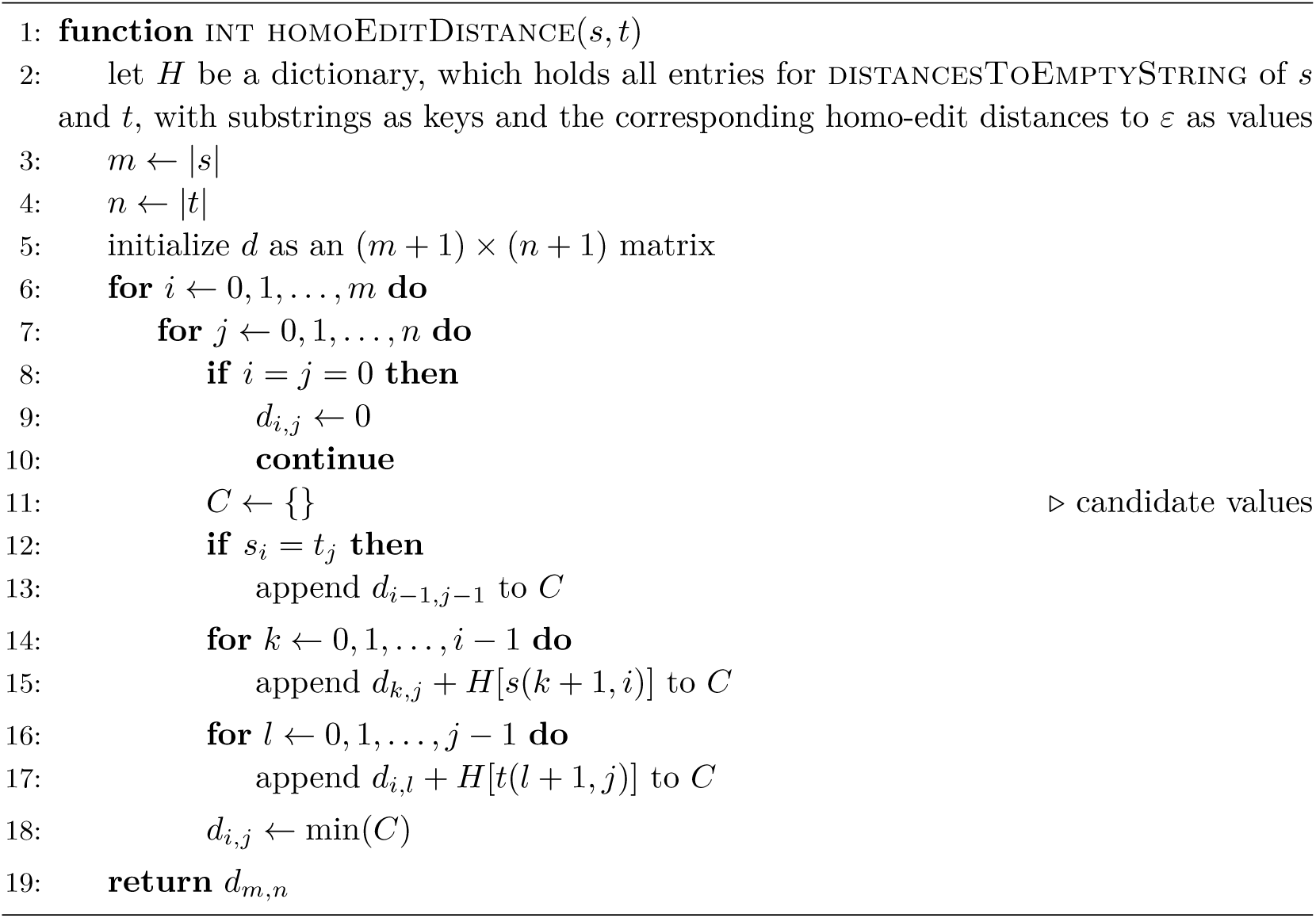

We now show how to precompute the homo-edit distances *H*(*s*(*i, j*), *ε*) between all substrings of a string *s* of length *n* and the empty string. Again, we use dynamic programming, filling an *n* × *n* matrix *h*(*s*), with the following recurrence:

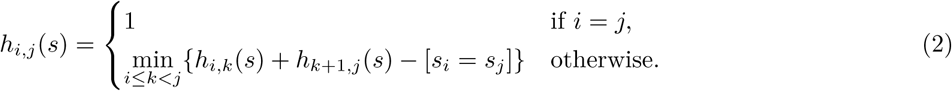

We start by initializing all homo-edit distances between *ε* and every substring *s*(*i, i*) of length one to *h*_*i,i*_(*s*) = *H*(*s*(*i, i*), *ε*) = 1 for all 1 ≤ *i*≤ *n*. Then we loop over all substrings of length two and compute their homo-edit distances to *ε*, and repeat the same procedure for all substrings of increasing length up to length *n*: To compute *h*_*i,j*_(*s*), we partition substring *s*(*i, j*) into all possible pairs of shorter substrings *s*(*i, k*) and *s*(*k* + 1, *j*), where *i* ≤ *k < j*. For each partition we compute the cost to delete it, and choose the minimum of these costs. If *s*_*i*_ ≠ *s*_*j*_ the cost of deleting a partition is the sum of the costs to delete either substring. If, however, *s*_*i*_ = *s*_*j*_ the cost decreases by one, which we notate using the Iverson bracket. The reason is that all partitions delete *s*_*i*_ in *s*(*i, k*) and *s*_*j*_ in *s*(*k* + 1, *j*) separately by two homo-deletions, but it is always possible to delete the characters at the first and last index together with one homo-deletion. In the end, *h*_*i,j*_(*s*) = *H*(*s*(*i, j*), *ε*) for all 1 ≤ *i < j*≤ *n*. See Algorithm 2 and the correctness proof below for more details.

#### Algorithm 2 Auxiliary dynamic programming algorithm to compute the homo-edit distance between every substring of a string *s* and the empty string.

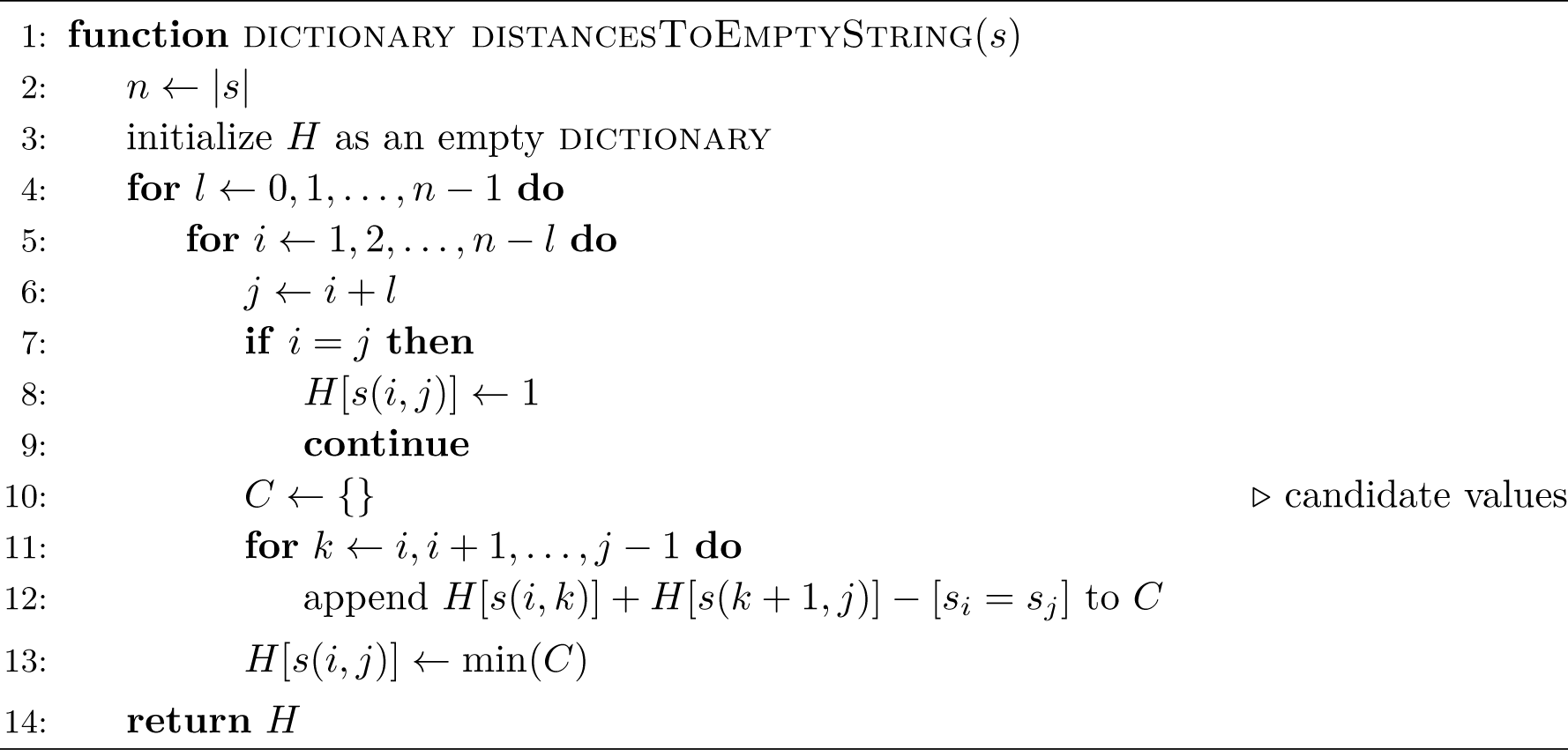

The example in Fig. 1 illustrates how the algorithms compute the homo-edit distance for the input strings *s* = `CTGCA` and *t* = `AGAAC`.

**Figure 1.**
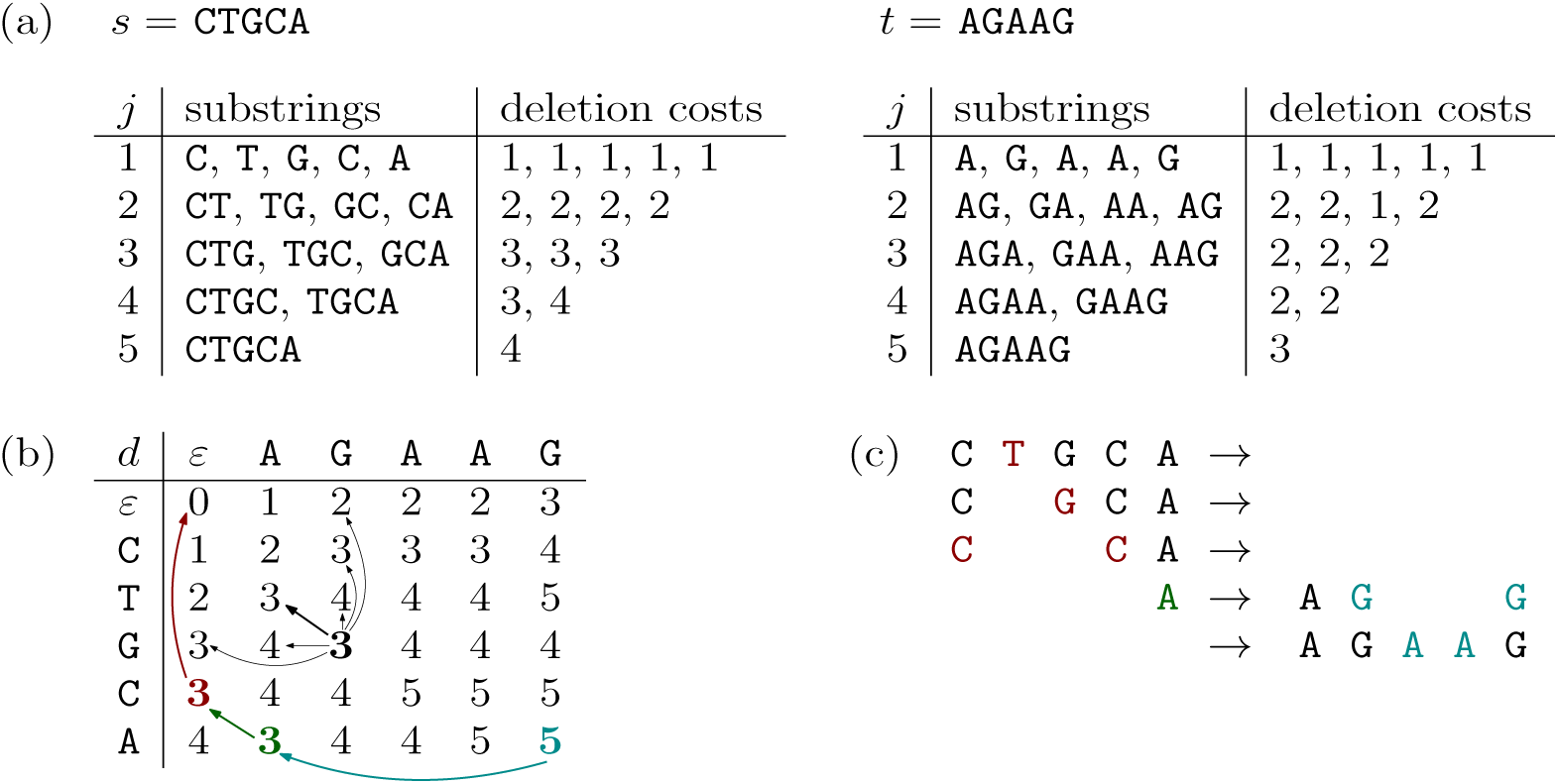
Example illustrating the algorithm. (a) Input strings *s* and *t* and the costs computed by the auxiliary DP. (b) Main DP matrix. An optimal backtracking path is colorized. Black arrows indicate how the entry for prefixes `CTG` and `AG` (shown in bold face) is computed. Here, the minimum is determined by the first case of Recurrence (1) and indicated by a bold arrow. (c) A transition in five steps as given by backtracking the colored path in (b).

### 4.2 Correctness

#### Lemma 4.

*Given a string s* = *s*_1_*s*_2_ … *s*_*n*_, *Recurrence* (2) *computes the homo-edit distance between every substring of s and the empty string ε.*

**Proof by induction.**

▪ *Base case: n* = 1. We need exactly one homo-deletion for one character, thus we have a homo-edit distance of 1, which is consistent with Recurrence (2).
▪ *Induction step: n* → *n* + 1. We consider two cases:
  ▪ Case 1: There exists an index *k* where 1 ≤ *k < n* + 1 such that *s*(1, *k*) and *s*(*k* + 1, *n* + 1) can be deleted independently from one another, i.e., we do not perform a homo-deletion that involves both substrings at once. This means the induction holds since we can reduce this problem to two subproblems.
  ▪ Case 2: There exists no such index *k*. Then *s*_1_ and *s*_*n*+1_ are the same character and must be deleted together because otherwise Case 1 would apply. That is, the cost for deleting *s*(1, *n* + 1) are the same as for *s*(1, *n*) because we can always delete *s*_1_ and *s*_*n*+1_ (and perhaps other equal characters in between) together with the last homo-deletion before reaching *ε*. ◂

#### Lemma 5.

*Given two strings s* = *s*_1_*s*_2_ … *s*_*n*_ *and t* = *t*_1_*t*_2_ … *t*_*m*_, *Recurrence* (1) *computes and stores the homo-edit distance between s and t in d*_*m,n*_.

**Proof.** The *edit distance problem* is to convert a string into another such that the sum of individual costs of the editing operations insertion, deletion, and substitution is minimized, where the mentioned editing operations can operate on exactly one character. Ukkonen [6] describes a generalization of this problem: Given two strings *s* = *s*_1_*s*_2_ … *s*_*n*_ and *t* = *t*_1_*t*_2_ … *t*_*m*_, we want to convert *s* into *t* such that the sum of individual costs of editing operations is minimized.

We can show that a problem is also a generalized edit distance problem by giving an *editing operation set E* ⊂ Σ^*^× Σ^*^, where an element (*x, y*), *x* ≠ *y*, represents an editing operation that replaces *x* with *y*, and a *recurrence* that defines a matrix *d* with *cost function δ* : *E* → ℕ as follows:

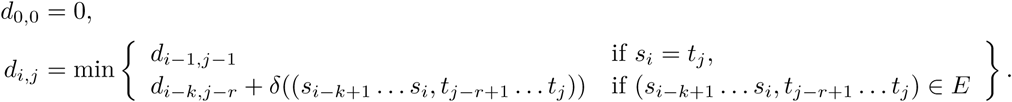

(Note that we rewrote Ukkonen’s recurrence to fit our notations.) Hence, if the homo-edit distance problem is a generalized edit distance problem, Recurrence (1) works correctly.

For the homo-edit distance problem we can represent the editing operation set as

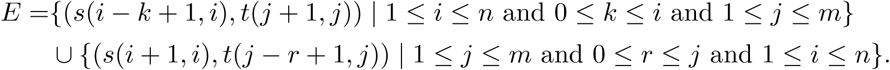

The cost function can be defined as

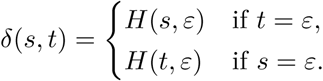

As a result, Algorithm 1 works correctly as we can rewrite Recurrence (1) as Ukkonen’s recurrence. ◂

### 4.3 Backtracking

From Lemma 1 and Lemma 3 we can deduce that an optimal series of operations needed for transforming *s* into *t* can be inferred from an optimal series of homo-deletions needed to transform both *s* and *t* into a common subsequence *v* with the property described in Lemma 3. Therefore, we disregard homo-insertions. Besides, we focus on backtracking one optimal series of homo-deletions that transform each input string into *v*. Note, however, that there might be multiple possible optimal series and subsequences.

In order to backtrack and thus generate an optimal series of homo-deletions as well as *v*, we augment our matrices *d* and *h* as follows: For each entry *d*_*i,j*_, we additionally store the indices of any entry *d*_*i*′,*j*′_ from which we came from. For each entry *h*_*i,j*_(*s*), we additionally store the smallest index *k* that led to *h*_*i,j*_(*s*). We proceed analogously for *h*(*t*).

Next, we backtrack a path from *d*_*m,n*_ to *d*_0,0_. Let 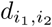 be the entry from where we obtained our current entry 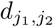. Let *v* be an empty string *ε* that will eventually hold our desired *v*, and let *L*_*s*_ and *L*_*t*_ be initially empty lists in which we will store our indices denoting an optimal series of homo-deletions from *s* or *t*, respectively. Note that homo-deletions cause indices to shift such that the indices stored in *L*_*s*_ and/or *L*_*t*_ might need to be adjusted accordingly. We consider three cases:

1. If we obtained 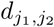 from a match, we prepend 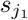 to *v*.
2. If we obtained 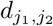 from an above entry that deletes *s*(*i*_1_, *j*_1_) from *s*, we recursively split the deletion of *s*(*i*_1_, *j*_1_) into the deletion of the two substrings *s*(*i*_1_, *k*) and *s*(*k* + 1, *j*_1_), where *k* is the respective index obtained from backtracking *h*(*s*). We abuse notation by using the same notation for any lower level of the recursion. The recursion adds a tuple (*k, k*) (or (*k* + 1, *k* + 1)) to *L*_*s*_ if a substring *s*(*k, k*) (or *s*(*k* + 1, *k* + 1)) consists of one character only. Every time we move up one recursion level, we check whether the outer characters 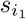 and *s*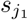 are equal. If so, from *L*_*s*_ we remove the tuple that is returned first by *s*(*i*_1_, *k*), which contains *i*_1_, as well as the tuple that is returned last by *s*(*k* + 1, *j*_1_), which contains *j*_1_. We then append (*i*_1_, *j*_1_) to *L*_*s*_.
3. If we obtained 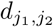 from a left entry that deletes *t*(*i*_2_, *j*_2_) from *t*, we proceed analogously to the second case.

### 4.4 Running Time Analysis

Consider *s* = *s*_1_*s*_2_ … *s*_*n*_ and *t* = *t*_1_*t*_2_ … *t*_*m*_. For string *s*, Algorithm 2 considers 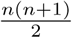 different substrings of *s*. For each of those substrings, we have up to *n* − 1 different partitions into two substrings. Consequently, the running time is in 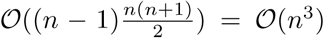. Analogously, we get 𝒪(*m* ^3^) to preprocess string *t*.

For each entry in *d*, Algorithm 1 regards up to *m* options from above, up to *n* options from left, and possibly one option from top-left. As we have *m* · *n* entries, the running time is in 𝒪 (*m* · *n* · (*m* + *n* + 1)) = 𝒪 (max *n, m* ^3^).

All in all, the running time is 𝒪 (*n*^3^) + 𝒪 (*m*^3^) + 𝒪 (max *{n, m}* ^3^) = 𝒪 (max *{n, m}* ^3^) and thus cubic in the input length. We can now state our main result:

#### Theorem 6.

*Algorithm 1 computes the homo-edit distance of two strings s* = *s*_1_*s*_2_ … *s*_*n*_ *and t* = *t*_1_*t*_2_ … *t*_*m*_ *in time 𝒪*(max*{n, m}*^3^).

**Proof.**

Follows from Lemmas 4 and 5 and the above running time analysis. ◂

## 5 Conclusions

The focus of this paper is to introduce the homo-edit distance problem and to present a solution to compute this distance in polynomial time. We have not yet considered applications of this distance to specific problems in bioinformatics and leave this as future work.

We can, for example, imagine applications to sequence analysis problems that involve tandem repeats, in a similar way as done by Sammeth and Stoye [5] who analyzed coding regions of the *Staphylococcus aureus* protein A gene (spa). *S. aureus* is a major human pathogen, and the analysis of relations between antibiotics-resistant strains can have important implications for clinical practice. Here, the homo-edit distance could be a good starting point for an all-against-all comparison of the spa-regions of different strains with an alphabet given by the repeats or higher order repeat structures.

Another possible application is the analysis of homopolymer-rich DNA-regions. Basecalling in these regions is particularly difficult for pyro- and ion torrent-based sequencing technologies, where over- and undercalling are common errors in these regions. The challenge is to distinguish these sequencing artifacts from true genetic content where a homo-edit distance-based analysis of the reads falling in such regions may provide some help.

In general, we envision also more theoretical work on extensions of the homo-edit distance. For which combinations of additional biologically meaningful operations like, e.g., duplications or mutations, can the distance still be computed in polynomial time and which versions become intractable? These and related open questions provide challenging opportunities for the theoretical bioinformatics community.

## Supplement Material

Source code available at https://github.com/AlBi-HHU/homo-edit-distance.

## Acknowledgements

The authors thank Max Ried for supporting the release of the code as a Python package.

## References

1 A. Apostolico and C. Guerra. The longest common subsequence problem revisited. Algorithmica, 2(1):315–336, 1987.

2 M. Brand. Das Homo-Edit-Distanz-Problem. Bachelor’s thesis, Heinrich Heine University Düsseldorf, 2020.

3 N. Jones and P. Pevzner. An Introduction to Bioinformatics Algorithms. MIT Press, Cambridge, MA, 2004.

4 S. Needleman and C. Wunsch. A general method applicable to the search for similarities in the amino acid sequence of two proteins. Journal of Molecular Biology, 48(3):443–453, 1970.

5 M. Sammeth and J. Stoye. Comparing tandem repeats with duplications and excisions of variable degree. IEEE/ACM Trans. Comput. Biol. Bioinformatics, 3(4):395–407, 2006.

6 E. Ukkonen. Algorithms for approximate string matching. Information and control, 64(1-3):100–118, 1985.

